# Reversible thiol oxidation increases mitochondrial electron transport complex enzyme activity but not respiration in cardiomyocytes from patients with end-stage heart failure

**DOI:** 10.1101/2022.05.27.493695

**Authors:** Ravi A. Kumar, Trace Thome, Omar Sharaf, Terence E. Ryan, George J. Arnaoutakis, Eric I. Jeng, Leonardo F. Ferreira

## Abstract

Cardiomyocyte dysfunction in patients with end-stage heart failure with reduced ejection fraction (HFrEF) stems from mitochondrial dysfunction that contributes to an energetic crisis. Mitochondrial dysfunction reportedly relates to increased markers of oxidative stress, but the impact of reversible thiol oxidation on myocardial mitochondrial function in patients with HFrEF has not been investigated. In the present study, we assessed mitochondrial function in ventricular biopsies from patients with end-stage HFrEF in the presence and absence of the thiol reducing agent dithiothreitol (DTT). Isolated mitochondria exposed to DTT had increased enzyme activity of complexes I (*p* = 0.009) and III (*p* = 0.018) of the electron transport system, while complexes II (p = 0.630) or IV (p = 0.926) showed no changes. However, increased enzyme activity did not carry over to measurements of mitochondrial respiration in permeabilized bundles. Oxidative phosphorylation conductance (*p* = 0.439), maximal respiration (*p* = 0.312), and ADP sensitivity (*p* = 0.514) were unchanged by 5 mM DTT treatment. These results indicate that mitochondrial function can be modulated through reversible thiol oxidation, but other components of mitochondrial energy transfer are rate limiting in end-stage HFrEF. Optimal therapies to normalize cardiac mitochondrial respiration in patients with end-stage HFrEF may benefit from interventions to reverse thiol oxidation that limits complex I and III activities.

## Introduction

Heart Failure with reduced Ejection Fraction (HFrEF) describes an inability of the heart to adequately pump blood due to impaired systolic function. End-stage HFrEF refers to a diagnostic category in which patients are non-responsive to conventional treatments and instead require advanced therapies like heart transplantation or placement of a mechanical pump to assist the heart (left ventricular assist device; LVAD). Insufficient ventricular function in HFrEF stems from contractile and energetic deficiencies. The mitochondria provide ~90% of ATP for continuously contracting cardiomyocytes, and myocardial mitochondrial function decreases with HFrEF [1–5]. Inadequate mitochondrial function leads to energy depletion, which contributes to contractile dysfunction, and correlates with disease severity [6] and patient mortality [7,8]. As such, targeting the mitochondria has gained increasing attention as an approach to improve myocardial function and disease outcomes in patients with HFrEF, especially for those that are not responsive to conventional pharmaceutical therapies [9–11].

One contributing factor to mitochondrial dysfunction is accumulation of reactive oxygen species (ROS) that results in excessive oxidation of proteins, lipids, and DNA. Cysteine residues are favorable targets for reversible redox reactions because of their thiol moiety (R-SH). Cysteine oxidation is a characteristic feature of normal cell signaling, however excessive oxidation can alter protein structure and function [12,13]. Mitochondria are considered a major source of increased oxidants in the failing heart and, by proximity, are also a primary target for redox reactions [12,13]. Redox proteomics in left ventricle biopsies from patients with end-stage HFrEF reveal increases in reversibly oxidized cysteine residues in proteins of complexes I and II of the electron transport system (ETS) in cardiac mitochondria [14]. Reversible oxidation of thiol moieties causes mitochondrial dysfunction in cardiomyocytes from mice fed a high fat, high sucrose diet [15] and from rats treated with doxorubicin [16]. Therefore, the mitochondrial limitations in patients with end-stage HFrEF may be caused to oxidative modifications of thiol redox switches that impair mitochondrial protein function.

The objective of this study was to determine if acute reduction of reversibly oxidized thiol moieties with dithiothreitol (DTT) could increase mitochondrial ETS complex activity and respiration at several levels of energetic demand in left ventricular biopsies from patients with end-stage HFrEF.

## Materials and Methods

### Human subjects

All experiments involving human subjects received approval by the Institutional Review Board of the University of Florida (protocol #202000853). Patients were informed of the nature and purpose of the study and signed a written consent in accordance with the Declaration of Helsinki. Patients with heart failure with reduced ejection fraction (HFrEF) undergoing placement of a left ventricular assist device (LVAD) were recruited to participate in this study. Our inclusion criteria included subjects 18 years or older with end stage HFrEF for more than one year. Patients with both ischemic and non-ischemic HFrEF were recruited. Patients were excluded from consideration if they presented with severe lung, hepatic or renal disease, or treatment with cancer chemotherapy.

### Biopsy Collection

Surgeries and biopsy collections were performed by cardiothoracic surgeons. As part of the LVAD placement, a portion of the left ventricle apical core was removed to insert the device. Apical core biopsies were immediately placed in ice-cold Buffer X (in mM: 7.23 K_2_EGTA, 2.77 Ca-K_2_EGTA, 20 imidazole, 20 taurine. 5.7 ATP, 14.3 PCr, 6.56 MgCl_2_-6H_2_O, 50 K-MES; pH 7.1). We rinsed and dissected myocardial tissue that was visibly free from fat, scarring, fibrosis and connective tissue, and immediately began prepping a portion of the sample for measurement of mitochondrial respiration. The remaining myocardial tissue was quickly flash frozen in liquid nitrogen for later isolation of mitochondria and measurement of ETS complex activity.

### Preparation of permeabilized cardiomyocyte bundles and mitochondrial respiration

Fine-tipped tweezers were used to separate cardiomyocyte bundles along the longitudinal axis to expose maximal cardiomyocyte surface area. We permeabilized ~2-4 mg (wet weight; WW) bundles in Buffer X containing saponin (30 µg/ml) for 30 min while gently rotating at 4°C. Bundles were then washed 3 × 5 min in Buffer D (in mM: 30 KCl, 10 KH_2_PO_4_, 5 MgCl_2_-6H_2_O, 105 K-MES, and 2.5 mg/ml BSA, 1 EGTA, pH 7.1). In bundles treated with DTT, the series of washes was done in the presence of 5 mM DTT in Buffer D. We chose this concentration based on a previous study [15]. We then proceeded with measurements of mitochondrial respiration.

We performed high-resolution O_2_ respirometry at 37°C in Buffer D supplemented with 5 mM creatine monohydrate and 10 µM Blebbistatin using an O2K Oxygraph (Oroboros, Innsbruck, Austria). To provide a comprehensive and physiologically relevant assessment of mitochondrial respiration, we implemented the previously described creatine kinase clamp that permits control over the extramitochondrial energy demand (ADP/ATP) [17,18]. Mitochondrial respiration has traditionally been measured under saturating conditions of ADP that overrides the mitochondrial internal mechanism of responding to energetic demand, whereas this approach permits assessment of mitochondrial oxygen consumption (*J*O_2_) at varying levels of energetic demand (ΔG_ATP_) that correspond to physiologically relevant levels of cellular ATPase activity. State 2 respiration was initiated by the addition of 10 mM pyruvate and 2 mM malate. The creatine kinase clamp was then established by the addition of 20 U/ml creatine kinase, 5 mM Tris-ATP and 1 mM phosphocreatine (PCr). These conditions (1 mM PCr) model near-maximal mitochondrial energetic demand *in vitro*. We then added 10 mM Cytochrome C to assess mitochondrial membrane integrity. Extramitochondrial energetic demand was then decreased by additional titrations of PCr (1-30 mM) to experimentally set different ATP free energy states through adjustments to the Cr/PCr ratio. Rate of oxygen consumption (*J*O_2_ in pmol/s) was calculated under each of these conditions and plotted against ΔG_ATP_ to establish a linear ‘force-flow’ relationship, where the slope represents the conductance of the electron transport system. Conductance measures the ability of mitochondria to respond to varying levels of energetic demand, akin to an exercise stress test. An overview of this assay, including a representative tracing of the rate of oxygen consumption and cellular conditions corresponding to different levels of PCr is shown in Figure 1 and Table 2. All measurements were conducted within an [O_2_] range of ~200 to 350 µM. Bundles were excluded from analysis if the addition of Cytochrome C resulted in a 15% increase in respiration, indicating mitochondrial outer-membrane damage during preparation. Following experiments, bundles were collected and freeze-dried using a bench top freeze dry system (Labconco, Kansas City, MO) and results were normalized by bundle dry weight (DW) (*J*O_2_ – pmol/s/mg DW). Where feasible, samples were run in duplicate, and results averaged together.

**Figure 1.**
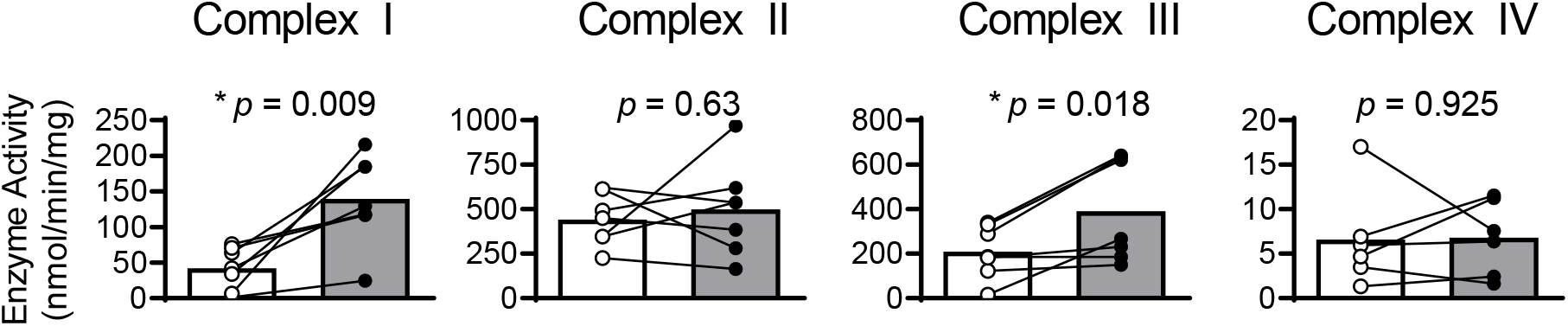
DTT treatment improves mitochondrial complex I and III enzyme activity. Enzyme activities of complexes of the electron transport system were assessed in isolated cardiomyocyte mitochondria from patients with HFrEF. 5 mM DTT treatment (grey) improved activity of complexes I and III compared to controls (white) but showed no effect on complexes II or IV. Enzyme activities compared via paired *t*-test. * *p* < 0.05.

We additionally calculated the corresponding [ADP] concentrations at each energetic state using the bioenergetics calculator provided by Fisher-Welman et al. [18]. These concentrations were used to determine mitochondrial ADP kinetics using the Michaelis-Menten equation to calculate maximal respiration (*J*O_2Max_ or V_max_) and the Michaelis constant (ADP_50_ or K_m_).

### Mitochondrial isolation

Mitochondria were isolated from previously frozen myocardial tissue using methods described in [19] with slight adaptation. Samples (~100-200 mg) were first incubated in PBS containing 10 mM EDTA and 3 mg/ml Collagenase Type I at 37 °C for 30 min to dissociate connective tissue in the especially fibrous human heart samples. All subsequent steps were performed on ice or at 4 ºC. Samples were minced for 5 minutes, and homogenates transferred to PBS with 10 mM EDTA and 0.5 mg/mL trypsin for an additional 5 min digestion. Next, we centrifuged samples at 500 x G for 5 min, poured off the supernatant, and resuspended the pellet in mitochondrial isolation medium (MIM; in mM: 50 MOPS, 100 KCl, 1 EGTA, 5 MgSO4) with 2 mg/mL fatty acid free BSA. Samples were then homogenized using a glass-teflon homogenizer (Wheaton) (6 passes at 40 rpm) and centrifuged at 800 × G to pellet nonmitochondrial myofibrillar proteins, nuclei, and other cellular components. The supernatant was collected and centrifuged at 10,000 × G for 10 min to pellet mitochondria, which were then gently resuspended in a small volume of MIM without BSA. We determined protein concentration of the resuspension using the DC Assay (BioRad) and diluted all samples to 2 mg/ml with MIM.

### Mitochondrial Complex Activity Assays

Mitochondrial complex activities were determined spectrophotometrically from isolated mitochondria using previously described methods [19,20]. Isolated mitochondria were diluted 1:1 in either hypotonic buffer (in mM: 25 K_2_HPO_4_, 5.3 MgCl, pH 7.2) for Complexes I and IV or Cell Lytic M (Millipore-Sigma) for Complexes II and III. Samples prepared in hypotonic buffer were subjected to four freeze-thaw cycles to optimize enzyme activity. All assays were run in duplicate under control conditions or in the presence of 5 mM DTT in a 96-well plate and changes in absorbance were tracked using a microplate reader (Synergy 2, BioTek/Agilent, Santa Clara, CA). This concentration was selected based on similar experiments in a previous study [15]. Background activities for each assay under control and DTT conditions were calculated during simultaneous reactions using inhibitors specific for each complex. Enzyme activity was calculated as the rate of change in absorbance after background subtraction.

Complex I (NADH:ubiquinone oxidoreductase) activity was measured in 10 mM KH_2_PO_4_ and 40 mM K_2_HPO4 (pH 7.5) with 3 mg/ml BSA, 240 µM KCN, 4 µM Antimycin A, 50 µM decyl-ubiquinone (DCU), and 80 µM 2,6-dichlorophenolindophenol (DCPIP) using 20 µg of mitochondria. We initiated the reaction with the addition of 1 mM NADH and monitored the reduction of DCPIP at 600 nm. Background activity assays were run in the presence of 25 µM rotenone. Changes in absorbance were monitored over 300 s and slopes were calculated between 100-200 s. Rates were converted to moles using the extinction coefficient 19100 M^−1^ cm^−1^ for DCPIP absorbance at 600 nm.

Complex II (succinate dehydrogenase; SDH) activity was measured in 10 mM KH_2_PO_4_, 2 mM EDTA and 1 mg/ml BSA (pH 7.8) supplemented with 4 µM rotenone, 0.2 mM Tris-ATP, 10 mM succinate, and 80 µM DCPIP using 5 µg of mitochondria. We initiated the reaction with the addition of 0.08 mM DCU and monitored the reduction of DCPIP at 600 nm. Background activity assays were run in the presence of 10 mM malonate. Changes in absorbance were monitored over 1200 s and slopes were calculated between 100-300 s for DTT-treated samples or 600-800 s for control samples. Rates were converted to moles using the extinction coefficient 19100 M^−1^ cm^−1^ for DCPIP absorbance at 600 nm.

Complex III (Q-Cytochrome C oxidoreductase) activity was measured in 10 mM KH_2_PO_4_, 2 mM EDTA and 1 mg/ml BSA (pH 7.8) supplemented with 0.2 mM Tris-ATP, 240 µM KCN, and 110 µM oxidized Cytochrome C using 15 µg of mitochondria. We initiated the reaction with the addition of 0.15 mM reduced decyl-ubiquinol (reduced) and monitored the reduction of Cytochrome C at 550 nm. Background activity assays were run in the presence of 10 µM myxothiazol. Changes in absorbance were monitored over 60 s and slopes were calculated between 15-45 s. Rates were converted to moles using the extinction coefficient 18500 M^−1^ cm^−1^ for Cytochrome C absorbance at 550 nm.

Complex IV (Cytochrome C Oxidase) activity was measured in 10 mM 10 mM KH_2_PO_4_, 250 mM sucrose and 1mg/ml BSA (pH 6.5) supplemented with 2.5 mM maltoside and 5 µM antimycin A using 5 µg of mitochondria. We initiated the reaction with the addition of 110 µM reduced Cytochrome C and monitored the oxidation of Cytochrome C at 550 nm. Background activity assays were run in the presence of 0.24 mM KCN. Changes in absorbance were monitored over 300 s and slopes were calculated between 120-180 s. Rates were converted to moles using the extinction coefficient 18500 M^−1^ cm^−1^ for Cytochrome C absorbance at 550 nm.

### Statistics

Data in text and tables are presented as mean ± SD. Individual data (scatterplots) and group means (bars) are shown in figures unless stated otherwise. We performed tests for normality (Shapiro-Wilk) before running paired *t* (parametric) or Wilcoxon signed rank (non-parametric) tests. We declared statistical significance when *p* < 0.05.

## Results

Patient characteristics are detailed in Table 1. Patients included in this study predominately exhibited HFrEF caused by non-ischemic (90%; n = 8) cardiomyopathy.

**Table 1.** Patient characteristics

|  | All Patients (n = 9) |
| --- | --- |
| Age (years) | 55.9 ± 14.0 |
| Male, % (n) | 77.8 (7) |
| EF (%) | 17.2 ± 8.3 |
| Ischemic CM, % (n) | 11.1 (1) |
| Hypertension, % (n) | 100 (9) |
| Hyperlipidemia, % (n) | 55.6 (5) |
| Atrial fibrillation, % (n) | 77.8 (7) |
| Asthma, % (n) | 11.1 (1) |
| COPD, % (n) | 22.2 (2) |
| CKD, % (n) | 55.6 (5) |
| Diabetes mellitus, % (n) | 44.4 (4) |
| CAD, % (n) | 55.6 (5) |

EF, ejection fraction; CAD, coronary artery disease; CKD, chronic kidney disease; CM, cardiomyopathy; COPD, chronic obstructive pulmonary disease.

Treating isolated mitochondria with DTT improved enzyme activity of complex I and III activity by 230% (*p* = 0.009) and 86% (*p* = 0.018), respectively, while having no significant effect on complex II (*p* = 0.630) or IV (*p* = 0.926) activity (Figure 1). An overview of creatine kinase clamp implemented to assess mitochondrial respiration, including a tracing of the rate of oxygen consumption and cellular conditions corresponding to different energetic states, is shown in Figure 2 and Table 2. Bundles treated with DTT displayed no significant differences in respiration under state 2 (*p* = 0.588) or highest energetic demand tested (1 mM PCr) (*p* = 0.312) conditions (Figure 3A). There were no significant differences in OXPHOS conductance (*p* = 0.439) between control and DTT-treated bundles (Figure 3). Mitochondrial ADP kinetics also did not differ between groups for maximal oxygen consumption (*J*O_2Max_) (*p* = 0.287) or ADP sensitivity (ADP_50_) (*p =* 0.514) (Figure 4).

**Table 2.** Bioenergetic conditions at each energetic state during the creatine-kinase clamp assay

| [PCr] (mM) | $\Delta G_{ATP}$ (kJ/mol) | [ADP] (mM) | [ATP] (mM) | [Cr] (mM) |
| --- | --- | --- | --- | --- |
| 1 | -54.16 | 0.176 | 4.82 | 4.824 |
| 3 | -57.09 | 0.058 | 4.94 | 4.942 |
| 6 | -58.32 | 0.037 | 4.96 | 4.963 |
| 15 | -60.63 | 0.016 | 4.98 | 4.984 |
| 30 | -62.37 | 0.009 | 4.99 | 4.991 |

Energetic conditions brought on by various titrations of PCr in the creatine-kinase assay. Concentrations determined by the bioenergetics calculator provided by Fisher-Wellman et al. (2018) [18].

**Figure 2.**
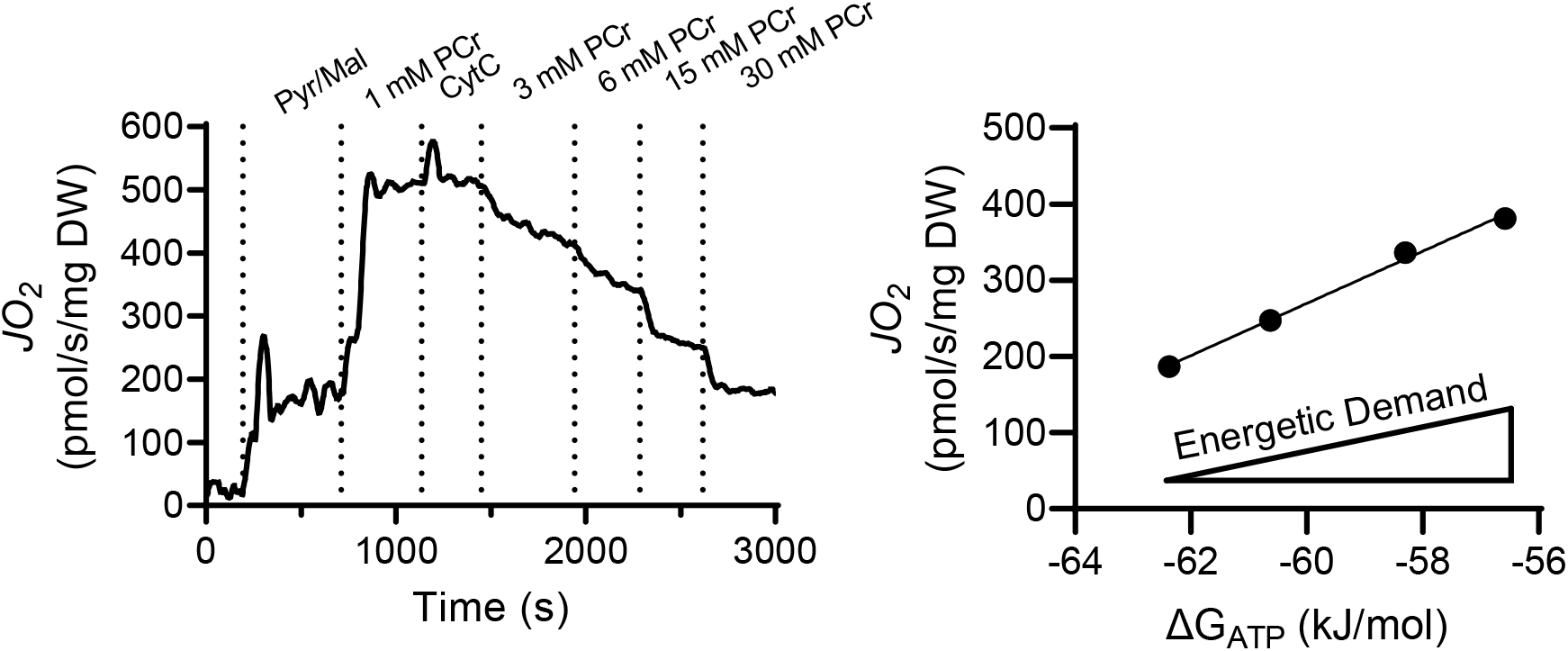
Creatine-Kinase Clamp to control mitochondrial energetic demand. Example tracing of the creatine-kinase clamp in which PCr is titrated to control the mitochondrial energetic demand (ΔG_ATP_) (see table 2). ΔG_ATP_ is then plotted against oxygen consumption (*J*O_2_) in which the slope of the relationship represents the mitochondrial oxidative phosphorylation (OXPHOS) conductance.

**Figure 3.**
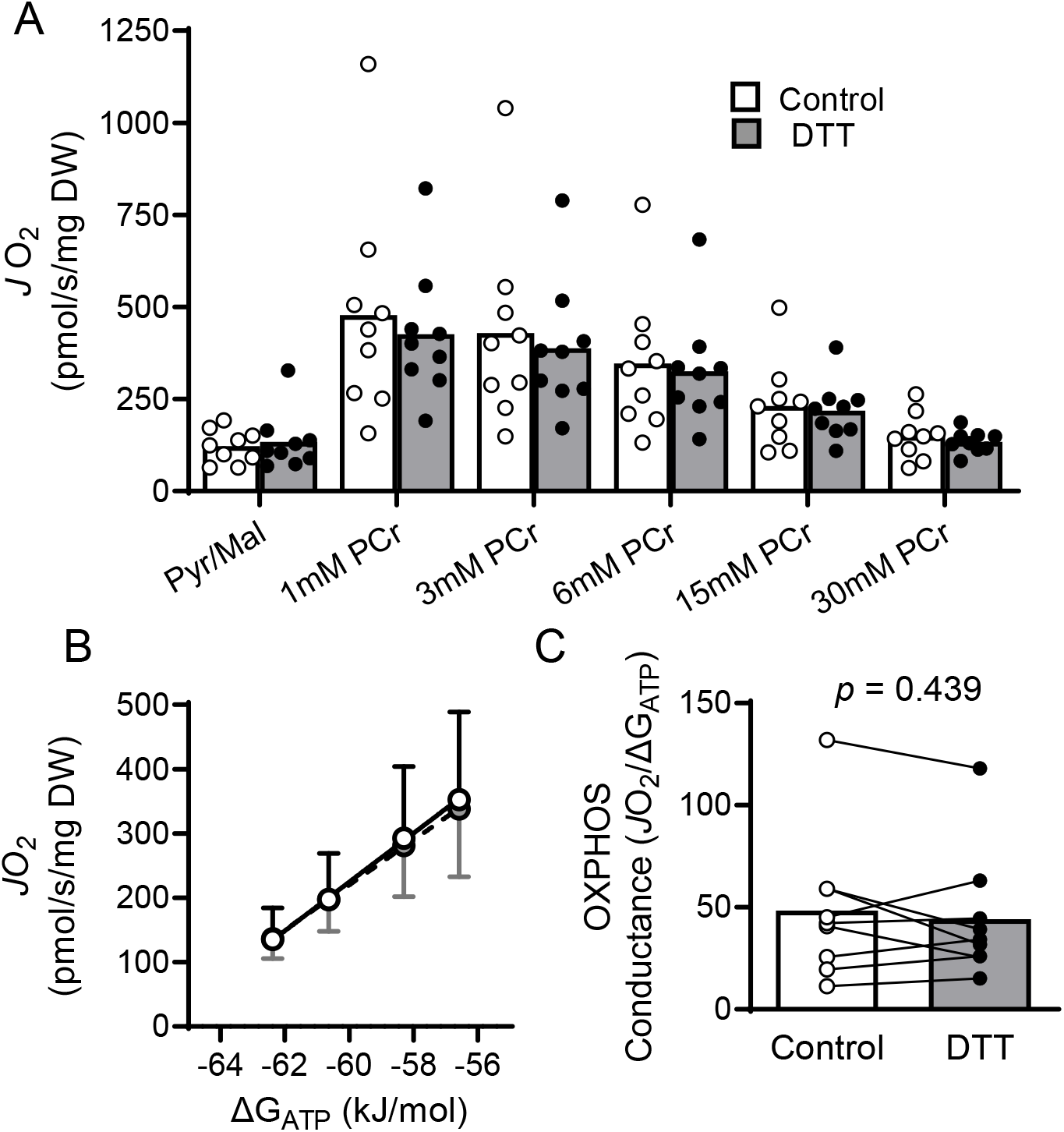
OXPHOS Conductance does not improve with DTT treatment. Saponin-permeabilized cardiomyocyte bundles from patients with HFrEF did not experience improvements in mitochondrial respiration under various energetic states following treatment with 5 mM DTT (A). Energetic demand (ΔG_ATP_) was manipulated through titrations of PCr (see Table 2) and plotted against the rate of oxygen consumption (*J*O_2_) (B) to determine OXPHOS Conductance (C), which was also unaffected by DTT treatment. *J*O_2_ and OXPHOS Conductance of control and DTT-treated samples compared via paired *t*-test.

**Figure 4.**
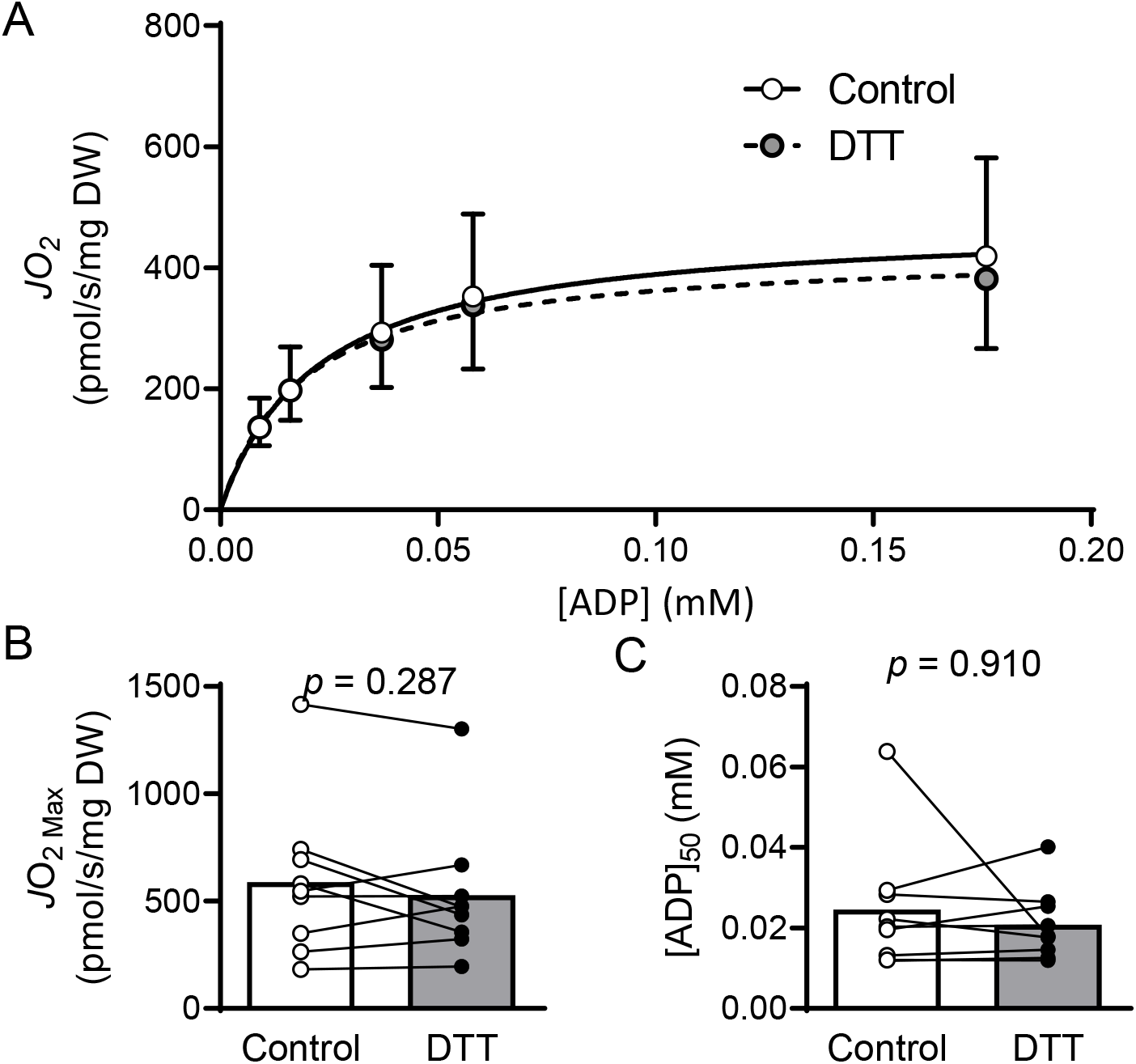
Mitochondrial ADP Kinetics are unchanged by DTT treatment. ADP concentrations corresponding to each energetic state were determined using the bioenergetics calculator from Fisher-Wellman et al. (2018) [18] (see Table 2) to determine Michaelis-Menton Kinetics and the effects of 5 mM DTT treatment from respiration experiments presented in Figure 2. DTT treatment had no effect on maximal respiration (*J*O_2Max_; compared via paired t-test) or ADP sensitivity ([ADP]50; compared via paired Wilcoxon signed rank test).

## Discussion

The main findings of this study are that the thiol-reducing agent DTT acutely increased the activity of mitochondrial ETS complexes I and III but did not change complex III and IV activities, mitochondrial oxidative phosphorylation conductance, or Michaelis-Menten kinetics of mitochondrial respiration in cardiomyocytes from patients with end-stage HFrEF. These findings suggest that reversible thiol oxidation, which impairs mitochondrial ETS complex activity, is not a limiting factor for mitochondrial metabolism and oxidative phosphorylation responses to carbohydrate-type substrates in the left ventricle of patients with end-stage heart failure.

### Mitochondrial bioenergetics and dysfunction in HFrEF

The heart is considered the most metabolically active tissue in the body owing to its continuous contractile activity. The lion’s share of this energetic demand is met by mitochondria [21], which occupy ~30% of cell volume within cardiomyocytes [22]. Severe and end-stage HFrEF involves a myocardial energetic crisis in which sufficient energy cannot be produced to meet the workload of the heart [1–5]. There is lower PCr/Cr ratio [7] and lower total adenine pool [23], as well as a shift in substrate preference from free fatty acid oxidation to glycolysis. However, impairments in mitochondrial respiration are also evident under controlled substrate conditions. The mitochondrial and energetic deficiency correlates with patient mortality and has been posited as a primary driving force of HFrEF disease pathology [8].

With greater recognition of the effect of mitochondrial dysfunction has on overall disease pathology, these organelles have become increasingly popular targets for potential therapeutics for patients with HFrEF [8–10]. Mitochondrial-targeted therapeutics may be especially valuable for patients with end-stage HFrEF where conventional treatments (e.g. ACE inhibitors and beta-blockers) are no longer effective. Investigations into myocardial mitochondrial function in ventricular biopsies from patients with end-stage HFrEF have revealed decreased mitochondrial respiration and activity of individual complexes of the electron transport system (ETS) [1–5]. In general, specific dysfunction within the mitochondria has been attributed to complex I [1–3,24,25], complex II [2,5], complex III [26], complex IV [2,25,27], and several enzymes within the TCA cycle [1,2,25].

The ETS couples the movement of electrons down a redox potential gradient with the active transport of protons (H^+^) against an electro-chemical gradient to generate a proton motive force that is then used to drive the formation high-energy phosphates in the form of ATP. Complexes I and II initiate the movement of electrons into the ETS by oxidizing reducing equivalents, while complex IV facilitates the final transfer of electrons to O_2_ to form H_2_O. Complexes I, III, and IV couple electron flow with proton pumping, while complex V uses the established proton motive force to phosphorylate ADP and regenerate molecules of ATP. The ratio of [ATP]/[ADP] is pushed far from equilibrium in favor of greater [ATP] by way of the proton motive force and it is the displacement from equilibrium that establishes the free energy associated with ATP hydrolysis (ΔG_ATP_) [28,29]. Flow of electrons through the ETS is limited by the development of the proton motive force which, in turn, is limited by the equilibrium of [ATP/[ADP]. As such, changes to [ADP] dictate the overall rate of mitochondrial respiration. [ADP] can change in response to increased workload or, in the case of the creatine kinase assay, by manipulating the following reaction: PCr + ADP ↔ Cr + ATP. Traditional approaches to measure mitochondrial function consist of measuring oxygen consumption under various energetic states that are induced by the addition of substrates, inhibitors, and high concentrations of ADP. While these experimental approaches have provided useful information about mitochondrial function, they disrupt the internal mitochondrial mechanisms of responding to energetic demand. Conversely, the mitochondrial creatine-kinase clamp permits control over mitochondrial energetic demand by titrating different concentrations of PCr to monitor mitochondrial oxygen consumption at various physiologically relevant energetic states [18,30,31].

The creatine-kinase clamp also provides several metrics that are similar to those presented with respiration experiments involving titrations of ADP up to saturating concentrations. State 2 refers to respiration in the presence of energizing substrates but in the absence of adenine molecules (ATP or ADP). Our values for state 2 respiration (Pyr/Mal) match those reported previously from patients with HFrEF when normalized by bundle wet weight (data not shown) [2]. Additionally, the energetic state elicited by 1 mM PCr represents a near-maximal energetic demand and is similar to the state 3 conditions elicited by maximal concentrations of ADP. The values of respiration at the highest level of energy demand within our conditions are similar to, or greater than, state III respiration presented in previous studies investigating myocardial mitochondrial function in patients with HFrEF [1,2,5]. These differences may be related to sample handling and preparation or experimental approach.

A traditional approach to assess mitochondrial respiration involves the use of Michaelis-Menten kinetics to define ADP sensitivity (K_m_; ADP_50_) and maximal capacity (V_max_) [5,32]. We calculated [ADP] at each energetic state using the energetics calculator provided by Fisher-Wellman et al [18] to determine Michaelis-Menten ADP kinetics of mitochondrial respiration. The K_m_ of cardiac mitochondria has been reported as 0.096 mM in healthy controls and 0.057 mM in patients with HFrEF [5]. These values reflect slightly lower [ADP] sensitivity than what we report here (0.025 mM in untreated samples), which likely results from differences in energetic conditions in the assay. Glutamate and malate were used as energizing substrates in the study cited above, while we used the combination of pyruvate and malate. Different substrate combinations can elicit differences in sensitivity and maximal respiration. Greater mitochondrial respiration in response to pyruvate over glutamate has been reported in murine hearts [18] and may carry over to sensitivity in humans. Additionally, when energetic states are controlled via titrations of PCr, maximal respiration is achieved at 0.176 mM of ADP, whereas traditional ADP titration experiments that disrupt the extramitochondrial energetic control expose mitochondria to upwards of 4 mM ADP. Acknowledgement of experimental bioenergetic conditions will be important to compare these results across studies. We also report *J*O_2 Max_ calculated from Michaelis-Menten ADP kinetics, which is essentially equivalent to *J*O_2_ at 1mM PCr discussed above.

### Myocardial oxidized shifts

A key feature of and contributing factor to myocardial dysfunction in HFrEF is excessive accumulation of reactive oxygen species (ROS). Markers of an oxidized shift in the cellular redox tone have been observed in ventricular biopsies from patients with HFrEF [14,25,33]. The mitochondria are a prominent source of increased ROS production and, by proximity, a target of redox reactions. While there are reports of decreased mitochondrial content that contribute to energetic insufficiency [5], Sheeran et al. [25] argue that mitochondrial dysfunction stems from oxidized post-translational modifications that impair enzyme and organelle function. They report increased markers of oxidation on subunits of complexes I, IV, and V of the ETS that correspond to decreased enzyme activity [25], while redox proteomics in ventricular biopsies also observed increased oxidation of subunits associated with complexes I and II [14]. Specifically, five subunits in Complex I showed increased markers of oxidation (carbonylation and nitrotyrosine) and all subunits contain Fe-S clusters that facilitate the transfer of electrons. Similarly, subunits I and II of complex IV contain redox centers that are directly involved in the transfer of electrons from cytochrome C to O_2_ [25]. These oxidative post-translational modifications have been suggested as contributors to mitochondrial enzymatic function and may be an important target to improve mitochondrial function and the energetic crisis in HFrEF [9,25].

Several processes within the mitochondria are regulated by redox switches which have been recently reviewed [12,13]. Specifically, several studies have investigated the role of thiol oxidation on mitochondrial function, predominately investigating the role of glutathiolynlation of cysteine residues [34–36]. Cysteines are commonly observed near active sites of enzymes because of their ability to stabilize Fe-S clusters. Two cysteines on the 75 kDa subunit of complex I (NDUSFI) (Cys^531^ and Cys^704^) are particularly sensitive to redox reactions that can impair enzyme function [37–39]. While specific cysteines were not investigated, the NDUSFI subunit experienced significant oxidative shifts in myocardial biopsies from patients with HFrEF compared to controls [14]. It follows that redox reactions on these residues may relate to impaired function that can be targeted with a reducing agent.

Dithiothreitol (DTT), also known as Cleland’s reagent, is a molecular compound characterized as a potent reducing agent in 1964 [40]. With its low redox potential (−0.33 V at pH 7), DTT is frequently used in molecular and biochemical experiments to directly reduce reversible oxidized disulfide bonds that form on cysteine. Acute DTT exposure *in vitro* has effectively improved enzyme activity following an oxidized shift in the cellular redox tone under several conditions. Improvements in striated muscle with DTT exposure include proteins involved in contractile [41] and mitochondrial function [15,16,42,43]. Mice on a high fat, high sucrose diet have decreased cardiac mitochondrial function due to impaired complex II activity that is attenuated with *in vitro* DTT treatment [15]. DTT treatment also improved mitochondrial dysfunction associated with drug toxicity in rats [16,44]. It’s important to acknowledge that direct exposure of isolated mitochondria to the thiol oxidizing agent diamide impaired complex I and III activity, and this was not attenuated with DTT treatment [37]. This may be due to different outcomes from reversible endogenous vs. irreversible exogenous oxidant exposure, or possibly through oxidant effects on the recruitment of binding partners that augment enzyme function [37].

Our study shows that reversible thiol oxidation also modulates mitochondrial ETS complex activity in the human heart. Specifically, our data provide evidence that DTT heightened enzymatic activity of complex I and III in mitochondria isolated from left ventricle of patients with end-stage heart failure. However, this increase in ETS complex activity did not translate into higher measures of mitochondrial respiration in permeabilized cardiomyocytes. The discrepant effect of DTT on activity of ETS complexes measures of respiration may be explained by several factors. First, enzyme activity assays were conducted in the presence of DTT, while respiration experiments consisted of a pre-treatment before measurements in the absence of DTT. It’s possible that thiol residues that were reduced during the DTT treatment during the wash stage were re-oxidized when cardiomyocytes were placed into the respiration buffer as cardiac mitochondrial ROS is increased in end-stage HFrEF. Second, protein thiols may have been more accessible for DTT in isolated mitochondria than in permeabilized bundles. Third, improved enzyme activity via reduction of oxidized thiol groups in Complex I and III may only detectable when the enzyme activity is not constrained by the mitochondrial membrane potential. As described above, the flow of electrons through the ETS in an in-tact system is limited at distinct control nodes, including establishment of the membrane potential [18,29]. Maximal activity of individual complexes may exceed that permitted by thermodynamic constraints of the complete system. Fourth, cardiac mitochondria complex I and III activities may not be rate limiting for respiration in patients with end-stage HFrEF. Other steps of mitochondrial energy transfer could limit respiration and oxidative phosphorylation conductance (e.g, matrix dehydrogenases, complex IV, and complex V/ATP synthesis). Our experiments found no improvement of Complex IV function with DTT. Additionally, cardiomyocytes of patients with end-stage HFrEF have lower abundance of CoQ10, which facilitates the transfer of electrons from Complexes I and II to complex III, that contributes to the mitochondrial dysfunction [45]. We did not measure the effects of DTT on the activity of matrix dehydrogenases or complex V as these assays are not justified by our results showing no effect of DTT on oxidative phosphorylation conductance.

## Limitations

There are several limitations to the current study. We did not have access to ventricular biopsies from control subjects to compare mitochondrial function from patients with HFrEF and the potential role of DTT. However, several previous studies have established decreased mitochondrial respiration in HFrEF using ventricular biopsies from patients with HFrEF compared to healthy donor hearts [1,2]. Additionally, DTT treatment had no effect [15] or even decreased [16] myocardial mitochondrial function in healthy rodent hearts, suggesting that any effects of DTT in our study would arise from reversing the oxidized thiol shift seen with end-stage HFrEF [25]. Our preparation focused on ETS complex activities, mitochondrial energy transfer conductance, and respiration under controlled conditions in the presence of glycolytic substrates. The shift in cardiac mitochondrial substrate in HFrEF is largely attributed to activation of a fetal-gene program that favors glycolytic over fatty acid metabolism [11], but matrix dehydrogenases associated with fatty acid oxidation may also undergo thiol oxidation that diminishes enzyme activity and contributes to the cardiac energy substrate shift in HFrEF. The substrate shift may be a protective compensatory response to an energy transfer system that is less ROS sensitive, as our study suggests to be the case for pyruvate metabolism by cardiac mitochondria. We did not determine which amino acid residues were affected by treatment with DTT. However, our goal was to examine the general impact of thiol oxidation on mitochondrial bioenergetics with the primary energy source in end-stage HFrEF. Finally, we did not have enough subjects to differentiate between effects in patients with ischemic and non-ischemic HFrEF. Mitochondrial dysfunction has been observed in both patient populations but is reportedly worse in ischemic HFrEF [2].

## Conclusions

Cardiac and mitochondrial redox imbalance are common features of HFrEF that contributes to dysfunction. Our study indicates that treatment with DTT, a potent reducing agent, can increase enzyme activity of Complexes I and III of the ETS in mitochondria isolated from HFrEF ventricular biopsies. These effects did not translate to greater mitochondrial respiration measured in permeabilized cardiomyocyte bundles, suggesting that other components of mitochondrial energy transfer are rate limiting for respiration and, presumably, ATP synthesis in end-stage HFrEF. However, our data indicate that human cardiac mitochondrial respiratory function can be modulated through reversible thiol oxidation. Therefore, optimal therapies to normalize cardiac mitochondrial respiration in patients with end-stage HFrEF will likely require interventions to reverse thiol oxidation that limits complex I and III activities in this population.

## Acknowledgments

The study was funded by the University of Florida (P0175597 - DRPD-ROF2020). R. Kumar received a predoctoral fellowship from the American Heart Association (20PRE35200047). L. Ferreira was supported by NIH R01 HL1331806 and University of Florida Research Foundation Professorship. T. Ryan was funded by NIH R01 HL149704. E. Jeng was supported by NIH R01 NR020175.

